# Adolescent Intermittent Ethanol Exposure Alters Adult Exploratory and Affective Behaviors, and Cerebellar *Grin2B* Expression in C57BL/6J Mice

**DOI:** 10.1101/2023.02.13.528396

**Authors:** Kati Healey, Renee C. Waters, Sherilynn G. Knight, Gabriela M. Wandling, Nzia I. Hall, Brooke N. Jones, Mariah J. Shobande, Jaela G. Melton, Subhash C. Pandey, H. Scott Swartzwelder, Antoniette M. Maldonado-Devincci

## Abstract

Binge drinking is one of the most common patterns (more than 90%) of alcohol consumption by young people. During adolescence, the brain undergoes maturational changes that influence behavioral control and affective behaviors, such as cerebellar brain volume and function in adulthood. We investigated long-term impacts of adolescent binge ethanol exposure on affective and exploratory behaviors and cerebellar gene expression in adult male and female mice. Further, the cerebellum is increasingly recognized as a brain region integrating a multitude of behaviors that span from the traditional primary sensory-motor to affective functions, such as anxiety and stress reactivity. Therefore, we investigated the persistent effects of adolescent intermittent ethanol (AIE) on exploratory and affective behaviors and began to elucidate the role of the cerebellum in these behaviors through excitatory signaling gene expression. We exposed C57BL/6J mice to AIE or air (control) vapor inhalation from postnatal day 28-42. After prolonged abstinence (>34 days), in young adulthood (PND 77+) we assessed behavior in the open field, light/dark, tail suspension, and forced swim stress tests to determine changes in affective behaviors including anxiety-like, depressive-like, and stress reactivity behavior. Excitatory signaling gene mRNA levels of fragile X messenger ribonucleoprotein (*FMR1)*, glutamate receptors (*Grin2a*, *Grin2B* and *Grm5*) and excitatory synaptic markers (PSD-95 and Eaat1) were measured in the cerebellum of adult control and AIE-exposed mice. AIE-exposed mice showed decreased exploratory behaviors in the open field test (OFT) where both sexes show reduced ambulation, however only females exhibited a reduction in rearing. Additionally, in the OFT, AIE-exposed females also exhibited increased anxiety-like behavior (entries to center zone). In the forced swim stress test, AIE-exposed male mice, but not females, spent less time immobile compared to their same-sex controls, indicative of sex-specific changes in stress reactivity. Male and female AIE-exposed mice showed increased *Grin2B* (Glutamate Ionotropic Receptor NMDA Type Subunit 2B) mRNA levels in the cerebellum compared to their same-sex controls. Together, these data show that adolescent binge-like ethanol exposure altered both exploratory and affective behaviors in a sex-specific manner and modified cerebellar *Grin2B* expression in adult mice. This indicates the cerebellum may serve as an important brain region that is susceptible to long-term molecular changes after AIE.

**Highlights:**

- Adolescent intermittent ethanol (AIE) exposure decreased exploratory behavior in adult male and female mice.
- In females, but not males, AIE increased anxiety-like behavior.
- In males, but not females, AIE reduced stress reactivity in adulthood.
- These findings indicate sex differences in the enduring effects of AIE on exploratory and affective behaviors.
- Cerebellar *Grin2B mRNA* levels were increased in adulthood in both male and female AIE-exposed mice.
- These findings add to the small, but growing literature on behavioral AIE effects in mice, and establish cerebellar excitatory synaptic gene expression as an enduring effect of adolescent ethanol exposure.

## 1. Introduction

Adolescence is a critical period for the development and maturation of neural circuitry, a process required for the emergence of cognitive and affective characteristics in adulthood (see (Larsen and Luna, 2018). Adolescent intermittent ethanol (AIE) exposure alters brain morphology and function in both humans and animal models (Brown et al., 2015; Crews et al., 2019), which may render individuals vulnerable to affective and cognitive dysregulation in adulthood (Dawson et al., 2008; Deykin et al., 1987). In humans, chronic alcohol use during adolescence is associated with persistent cognitive and affective deficits (Spear, 2018), as well as increased risk of developing alcohol use disorder in adulthood (Dawson et al., 2008). Studies in animal models using rats show persistent effects of AIE on some affective-like behaviors including ethanol drinking, anxiety-like behavior, impulsivity, behavioral flexibility, memory, sleep, and social anxiety (Crews et al., 2019; Dannenhoffer et al., 2018; Healey et al., 2022; Pandey et al., 2015; Robinson et al., 2021). Thus, the data from clinical, and some basic research studies, support the conclusion that binge-like ethanol exposure during adolescence creates a propensity toward affective dysregulation (Dannenhoffer et al., 2018; Gilpin, 2012; Healey et al., 2022; Pandey et al., 2015; Varlinskaya et al., 2017). However, to better understand the neurobehavioral mechanisms underlying persistent AIE effects on affective behavior, a more fine-grained analysis of the specific nature of AIE effects on several affective behaviors in multiple preclinical models is needed.

Sex differences in the acute and long-term effects of adolescent ethanol exposure are increasingly found in clinical and preclinical research. In rats, AIE exposure sex-specifically alters cognitive flexibility (Macht et al., 2020), affective behaviors (Healey et al., 2022; Varlinskaya et al., 2020), fear conditioning (Chandler et al., 2022), and social drinking (Towner et al., 2022) in adulthood. We recently showed that male and female AIE-exposed mice showed sex-specific patterns of subsequent ethanol drinking and withdrawal period-dependent differences in affective behaviors.

Specifically, in C57BL/6J mice anxiety-like behavior (center zone distance traveled) was assessed following experimenter-administered AIE followed by short or protracted withdrawal and subsequent voluntary ethanol alcohol drinking and anxiety-like behavior measures using the open field test (OFT). AIE males had increased ethanol preference compared to male control animals at both withdrawal periods, however AIE treated females exhibited reduced ethanol preference compared to controls only with the protracted withdrawal period. Further, females were more susceptible to short-term withdrawal and subsequent voluntary drinking to reduce the distance traveled in the OFT and in the center zone (anxiety-like behavior), whereas males modest reductions in center zone activity only manifested at the protracted withdrawal prior to voluntary drinking (Maldonado-Devincci et al., 2022). Other studies found sex differences in C57BL/6J mice that were treated with AIE and then received restraint stress in adulthood, with females showing greater AIE and stress-induced novelty-induced hypophagia compared to males (Kasten et al., 2020). Together, these data indicate that AIE alters adult behavioral characteristics, some of which are sex-specific, but more work is needed to understand the underlying long-lasting neurobiological changes that mediate these behavioral alterations.

The relationship between cerebellar and affective functioning is increasingly appreciated in the clinical literature, indicating cerebellar involvement in disorders such as attention deficit disorder, schizophrenia, bipolar disorder, depression, anxiety disorders and importantly, cerebellar dysfunction is the mechanism underlying the cerebellar cognitive affective syndrome (Baek et al., 2022; Depping et al., 2018; Ke et al., 2016; Liao et al., 2010; Phillips et al., 2015; Schmahmann and Sherman, 1998). Several of these disorders have an anxiety component, although this may be bidirectional and topographically mediated (Chin and Augustine, 2023). In rodent models, lesions of the cerebellar vermis have been found to increase anxiety-like behavior (Bobée et al., 2000; Supple et al., 1987), and other areas have no effect on anxiety behaviors indicating that not all cerebellar cortices influence anxiety behaviors (for review, see (Chin and Augustine, 2023). Models of either purkinje cell neurodegeneration (Spinocerebellar ataxia rodent model) or total purkinje cell loss (Lurcher mouse) in the cerebellum have largely been found to exhibit anxiolytic behavior (Boy et al., 2009; Hilber et al., 2004; Lorivel et al., 2010; Monnier and Lalonde, 1995); however, some anxiogenic behaviors have been reported (Kelp et al., 2013). Together, these clinical and pre-clinical data support a complex role of the cerebellum in anxiety behaviors and clinical disorders with an anxiety component that warrants further investigation. Additionally, it is notable that the cerebellum is highly sensitive to the effects of acute and chronic ethanol exposure (Coleman et al., 2014; Dong et al., 2022; Meruelo, 2020; Sullivan et al., 2020) though the possible enduring impacts of adolescent ethanol exposure on the cerebellum (Rapp et al., 2022) are only beginning to be addressed. For this reason, we focused the biological measures in this series of experiments on the cerebellum, rather than on the hippocampus, which has been more thoroughly studied in the context of AIE. Furthermore, our previous work has shown sex-specific changes in ethanol drinking and behavior following AIE exposure and voluntary access to ethanol in adulthood (Maldonado-Devincci et al. 2022). The present study extended these studies to examine the sex-specific impacts of AIE on behavior in adulthood in mice that did not have a history with ethanol drinking after AIE exposure.

Of note, affective regulation is also mediated through bidirectional connectivity between the hippocampus and the cerebellum (see (Yu and Krook-Magnuson, 2015) for review); as are spatial and temporal information processing (Fries, 2005; Onuki et al., 2015; Rochefort et al., 2011), behavioral conditioning (Takehara et al., 2003, 2002), and epileptogenesis (Krook-Magnuson et al., 2014). We and others have shown the hippocampus to be highly sensitive to the effects of AIE on both morphological (Mulholland et al., 2018; Risher et al., 2015a, 2015b; Swartzwelder et al., 2020) and physiological (Fleming et al., 2013, 2012; Li et al., 2013; Risher et al., 2015a, 2015b; Swartzwelder et al., 2017) endpoints. At a physiological level, there is a functional coupling of hippocampal and cerebellar neural network activity under certain behavioral circumstances that is manifested as a synchronization of theta oscillation activity across the two structures (Wikgren et al., 2010). This indicates ongoing synaptic co-regulation between the cerebellum and hippocampus that may be significant for both cognitive and affective functioning – both of which are altered by AIE (see (Crews et al., 2019; Robinson et al., 2021).

Recently, it was shown that expression of various genes in the cerebellum (e.g., Fragile X Messenger Ribonucleoprotein 1 (*Fmr1*) and its interactive genes, postsynaptic density 95 (*Psd95*), excitatory amino acid transporter 1(*Eaat1*), subtype of metabotropic glutamate receptor (*Grm5*), and ionotropic glutamate N-methyl D-aspartate (NMDA) receptor subunits (*Grin2B)* are altered by either acute or chronic ethanol exposure in adult rats (Dulman et al., 2021, 2019). However, the effects of adolescent ethanol exposure on synaptic genes particularly *Fmr1* and related genes in the cerebellum in adulthood are not known. To address these gaps, the present experiments were designed to explore the effects of AIE on affective behavior in adult male and female mice (which have been used far less than rats in rodent studies of AIE) more fully and to provide an initial assessment of AIE on cerebellar synaptic-relevant gene expression.

## 2. Methods

### 2.1 Subjects and Ethanol Exposure

Adolescent male and female C57BL/6J mice (n=8/group/sex) were obtained from Jackson Laboratories (Bar Harbor, ME) on PND 21. All mice were allowed to acclimate to the colony for one week (PND 21-27) prior to experimentation, where they were handled daily (animals were picked up, oriented to the injection position and weighed) to allow for acclimation to experimenter manipulation. Mice were PND 28 at the beginning of the experiment. Animals were group-housed (4-5 per cage) throughout the experiment, with free access to food and water. A visual depiction of the experimental procedure is shown in **Figure 1**. Food and water were available *ad libitum* throughout the experiment. All mice were maintained in a temperature and humidity-controlled room with lights on from 0700-1900 hr. Animal care followed National Institutes of Health Guidelines under North Carolina Agricultural and Technical State University Institutional Animal Care and Use Committee approved protocols.

**Figure 1:**
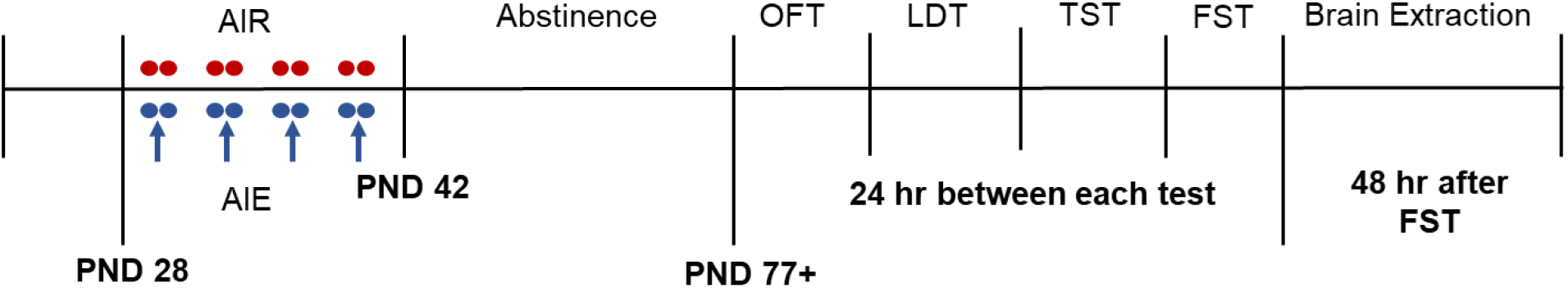
Experimental Timeline. Mice were exposed to AIE or AIR between PND 28-42 followed by long-term abstinence. Mice were then run over four consecutive days between PND 77-86 in the open field test (OFT), light/dark test (LDT), tail suspension test (TST), and forced swim stress test (FST). Twenty-four hours elapsed between each behavioral test. Brains were extracted 48 hours after the forced swim stress test. Red circles indicate air exposure. Blue circles indicate adolescent intermittent ethanol (AIE) exposure. The blue arrows indicate blood ethanol concentration (BEC) assessment.

Mice were exposed to repeated intermittent air or ethanol vapor using previously established protocols (Maldonado-Devincci et al., 2022). Briefly, mice were exposed to air (controls) or ethanol (experimental) vapor inhalation using four two-day ON/ two day OFF exposure cycles from PND 28-42. One cycle is defined as two days of vapor inhalation exposure (ON: air or ethanol) followed by two days of no exposure (OFF). Blood ethanol concentrations were stabilized using the alcohol dehydrogenase inhibitor pyrazole (1 mmol/kg, as used in previous work (Becker and Lopez, 2004; Jury et al., 2017; Maldonado-Devincci et al., 2022, 2016, 2014b). On each exposure day, mice were weighed and administered an intraperitoneal injection (0.02 ml/g) of pyrazole combined with saline for control mice or combined with 1.6 g/kg ethanol (8% w/v) for ethanol-exposed mice and immediately placed in the inhalation chambers (23 in x 23 in x 13 in; Plas Labs, Lansing, MI) for 16 hr overnight with room air (control group) or volatilized ethanol (ethanol group) delivered to the chambers at a rate of 10 liters/min. Ethanol (95%; Pharmco by Greenfield Global) was volatilized by passing air through an air stone (gas diffuser; Saint Gobain) submerged in ethanol.

On PND 29, 33, 37, and 41, at 0900 hr, mice were removed from the vapor inhalation chamber and 50 μL of blood was collected from the submandibular space for blood ethanol concentration (BEC) assessment and then returned to the home cage for 8 hr. On intervening days, mice were left undisturbed in the home cage. All blood samples were centrifuged at 5,000*g* and serum was collected and used to analyze BECs using the AM1 blood alcohol analyzer (Analox Instruments, Lunenburg, MA). Mice were left undisturbed in the home cage, except for regular cage maintenance until behavioral testing began. Behavioral testing began in adulthood at PND 77 or later, an abstinence period of 34 days or greater.

### 2.2 Behavioral Testing

Each mouse underwent four consecutive days (between PND 77-86) of behavioral testing investigating exploratory and affective behaviors using the (1) open field test (OFT), (2) light/dark test (LDT), (3) tail suspension test (TST), and (4) forced swim stress test (FST). In the OFT, distance traveled (cm), rearing (frequency), time spent in the center zone (seconds), entries into the center zone (frequency), and distance traveled in the center zone (cm) were quantified. In the LDT, latency to enter the light side (seconds), time on the light side (seconds), and entries into the light side (frequency) were quantified. In the TST, latency to immobility (seconds) and time immobile (seconds) were quantified. In the FST, time immobile (seconds) and struggling behavior (seconds) were quantified. All mice were counterbalanced across days and across groups for testing. Mice were transported and acclimated to the behavioral testing room for at least 60 min before behavioral testing. For each behavioral test, the testing arena was cleaned with 70% ethanol and allowed to dry completely before the next mouse was introduced. All behavioral tests were video recorded for offline analysis with the experimenter blind to sex and exposure conditions. Experimental details are described below.

#### 2.2.1 Open Field Test

Mice were tested for a 60 min trial in the OFT (Tatem et al., 2014). After acclimation, mice were introduced to the OFT under regular overhead lighting (400 lux). Mice were introduced to the testing arena (40.6 cm x 40.6 cm) to assess general exploratory behaviors (distance traveled, rearing) and anxiety-like behavior (center zone activity). Immediately upon removal from the arena, mice were returned to their home cage and to the colony. Trials were video recorded and data were quantified offline using AnyMaze software (version 7.0).

#### 2.2.2 Light/Dark Test

The LDT is used to evaluate anxiolytic behavior in rodents as they have a natural aversion to brightly lit environments (Takao and Miyakawa, 2006). With this test, anxiety in mice can be measured through motor activity and time spent in the light compared to dark compartments of the apparatus (Himanshu et al., 2020). This test was conducted within the same chamber as the OFT test with an opaque divider (20.3 cm x 20.3 cm) inserted to create the dark compartment. After acclimation mice were placed in the dark compartment for one min before the 10 min trial began. The light side of the box was illuminated with a bright light (850 lux). Videos were scored by experimenters blind to experimental conditions for latency to enter light side, duration of time on light side, and entries to light side.

#### 2.2.3 Tail Suspension Test

The tail suspension test (TST) attempts to translate depressive-like behaviors in humans by observing initial escape-oriented movements made when the rodent is placed in an inescapable environment (Cryan et al., 2005). The TST was conducted using previously established procedures (Can et al., 2012). After acclimation mice were briefly restrained to pass the tail through a small plastic cylinder to prevent tail climbing and then attach an adhesive strip to the end of the tail. Mice were then suspended 60 cm above the ground for a duration of 6 min. Mice were then immediately returned to their home cage after the adhesive strip and plastic cylinder were removed. Sessions were recorded and scored offline by experimenters blind to experimental conditions for latency to immobility and duration of immobility.

#### 2.2.4 Forced Swim Stress Test

Mice were tested for response to the FST using previously established protocols (Maldonado-Devincci et al., 2016, 2014a). Glass cylinders (Wholesale Flowers and Supplies; 20 cm diameter, 40 cm height) were used as the swim test apparatus. The glass cylinders were filled with water (23–25°C) to approximately 20 cm in depth and were separated by a visual barrier. After acclimation, mice were removed from the home cage and gently lowered into the apparatus for a 10 min swim session. Mice were removed, lightly dried, and returned to the home cage and placed on a warming pad for 15–20 min. Latency to immobility, time immobile, and struggling/climbing were assessed by experimenter-manual scoring blind to experimental conditions after the session.

Immobility was defined as the mouse remaining completely still or with minimal involuntary movements required to remain afloat. Struggling/climbing was defined as vigorous swimming, thrashing, and/or climbing in an attempt to escape, which usually subsided by the first three minutes after presentation to the inescapable swim tank; therefore, this behavior was only scored for the first three min of the session. After each swim session, the cylinders were emptied, cleaned, and refilled with fresh water for the next subject.

### 2.3 Brain Tissue Collection

Forty-eight hours following the FST, brains were collected. Mice were euthanized by rapid decapitation, brains were extracted, and tissue was harvested for various brain regions, including the cerebellum. Brains were snap frozen and stored at -80⁰C until analysis. Target genes were fragile X messenger ribonucleoprotein (FMR1), excitatory amino acid transporter 1 (Eaat1/Slc1a3), glutamate metabotropic receptor 5 (Grm5), glutamate ionotropic receptor NMDA type subunit 2A (Grin2A) and 2B (Grin2B), postsynaptic density marker 95 (PSD-95, Dlg4), and Hypoxanthine phosphoribosyltransferase 1 (Hprt, housekeeper mRNA).

### 2.4 RNA isolation and measurement of mRNA levels

Whole, mouse cerebellar tissue was crushed on dry ice with a mortar and pestle, and total RNA was extracted and purified from an aliquot of the tissue using the miRNeasy Mini Kit protocol and DNase Kit (Qiagen). A mixture of all subregions were included in the tissue aliquot used for RNA isolation. RNA was reverse transcribed to cDNA using the High-Capacity cDNA Reverse Transcription Kit (Applied Biosystems). cDNA was amplified in duplicate on a CFX Connect qPCR system (Bio-Rad) using PowerUp SYBR master mix (Applied Biosystems). cDNA was amplified using primers for target genes (**Table 1**) with *Hprt*, hypoxanthine–guanine phosphoribosyltransferase, used as a reference gene. Using the ΔΔCt method (Livak and Schmittgen, 2001), relative mRNA levels of target genes were normalized by subtracting the Ct value of *hprt* from the Ct value of the target gene, and delta Ct values for all samples were then normalized to the mean of the control group. Results are represented as the fold change in mRNA levels of the ethanol treated group compared to the control group.

**Table 1:**
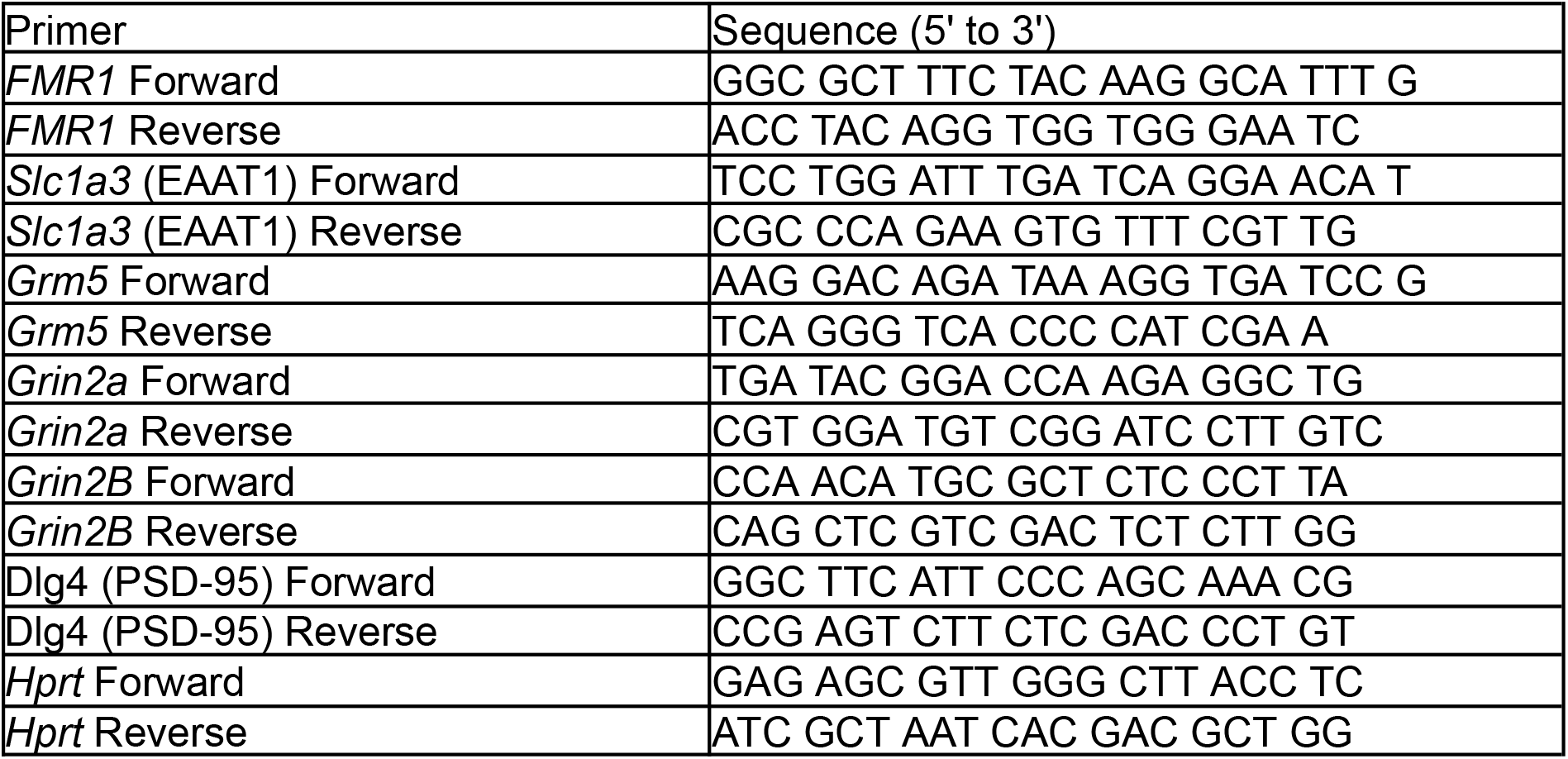
Various Primers used to measure mRNA levels.

### 2.5 Statistical Analyses

All behavioral data were analyzed using a two factor between subjects ANOVA with Exposure (2; Air, Ethanol) and Sex (2; Male, Female) as factors. All videos were quantified with the experimenter blind to sex and exposure. In the presence of significant interactions, Sidak’s multiple comparisons post hoc tests were used where appropriate. For each gene, fold changes within each group were assessed for normality. Student’s t-tests were performed for all normally distributed data. Fold change results within the female AIE group Grin2a mRNA results did not pass the Kolmogorov-Smirnov normality test, so a Mann-Whitney Rank Sum test was conducted. The mRNA data were analyzed separately for each sex for each target using a student’s t-test.

## 3. Results

### 3.1 Blood ethanol concentrations

Blood ethanol concentrations (BECs (mg/dl); **Table 2**) varied across cycles (F_(3,42)_ = 38.92, *p<0.0001*). Post hoc analyses for BECs collapsed across sex indicate that cycles 1 and 3 were lower than cycles 2 and 4, and cycle 3 was lower than cycle 1. Although the main effect of Sex achieved statistical significance (F_(1,14)_ = 5.29, *p=0.0374*), post hoc analyses did not reveal any statistically significant differences between males and females at any given cycle. There were no significant correlations between BECs and distance traveled. Males r=0.05, p = 0.93; Females r=0.54, p = 0.17.

**Table 2:**
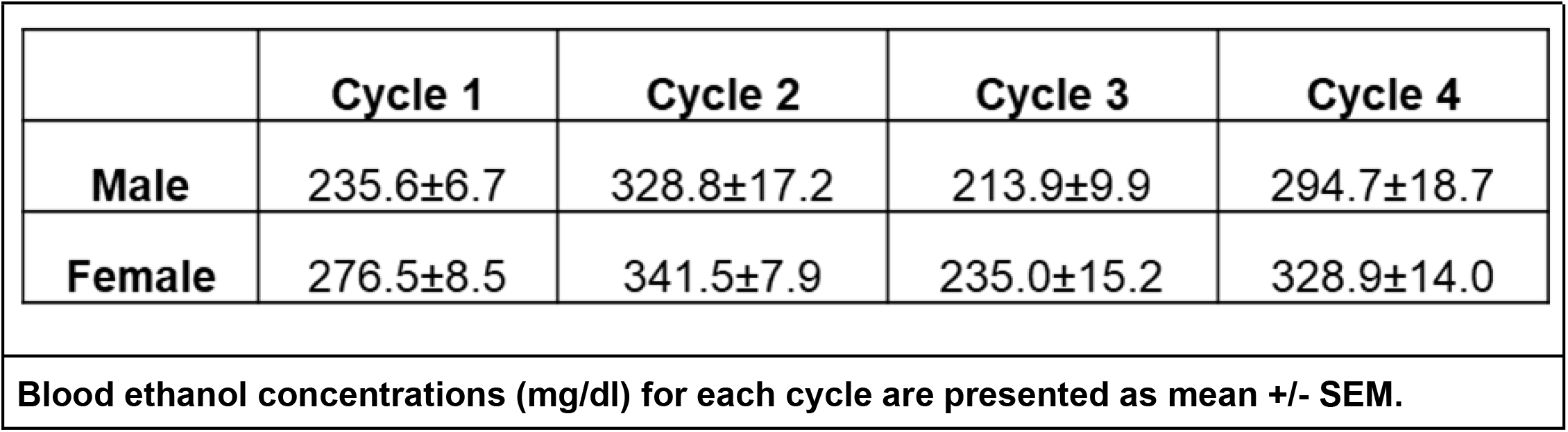
Blood Ethanol Concentrations.

### 3.2 Behavioral Testing

All mice underwent a series of behavioral tests over four consecutive days. This included the OFT, LDT, TST, and FST to determine changes in exploratory behavior, anxiety-like, depressive-like behavior, and stress reactivity, respectively. All statistical analyses are presented in **Supplemental Table 1**.

#### 3.2.1 Open Field Test

To assess overall activity in the open field, we measured the total distance moved in the apparatus during the trial. AIE reduced the total distance moved in both male and female mice (Main effect of Exposure: F_(1,28)_ = 31.49, *p<0.0001*) (**Fig. 2A**). In addition, AIE reduced the frequency of exploratory rearing in the open field in female mice (Exposure x Sex interaction: F_(1,28)_ = 4.71, *p=0.0385*; Main effect of Exposure: F_(1,28)_ = 9.26, *p=0.0044*) (**Fig. 2B**).

**Figure 2:**
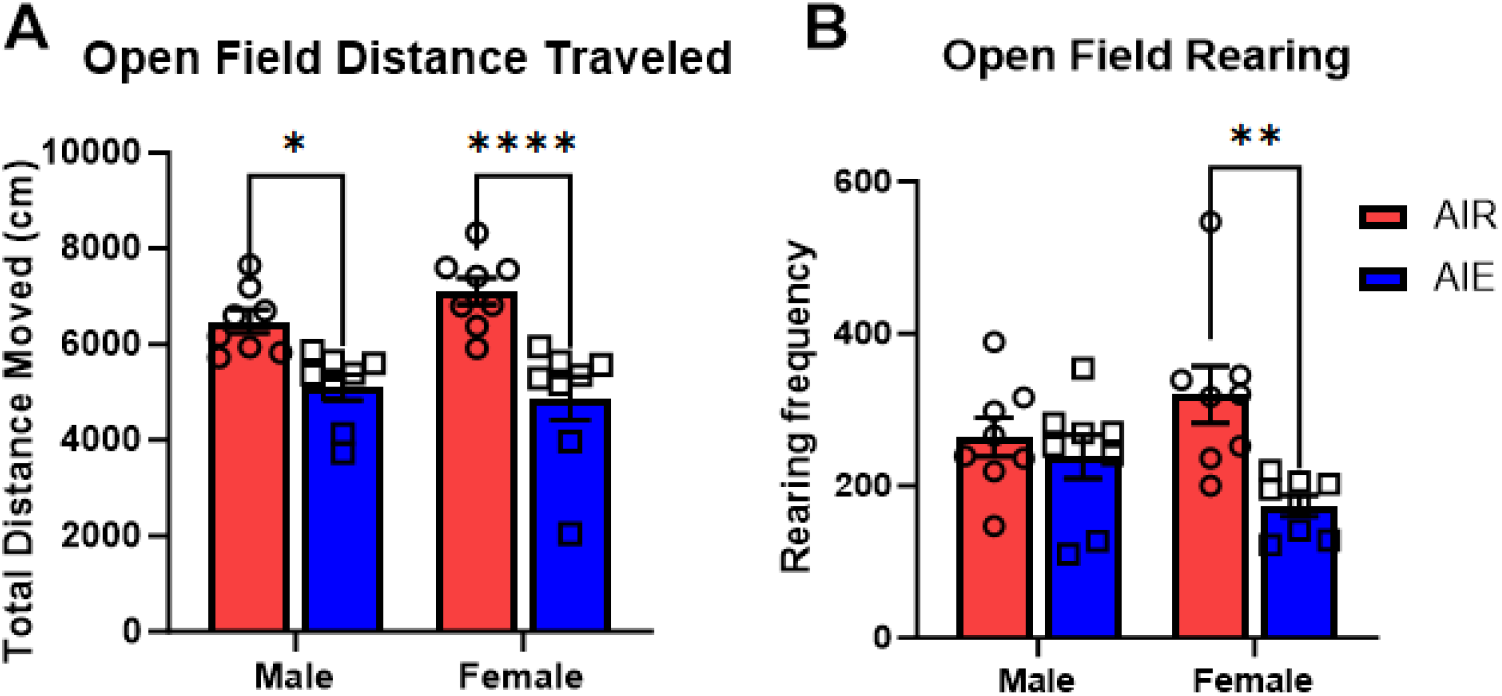
Open Field Data. A) Total distance traveled (cm) in the entire arena, B) Rearing Frequency. Data are presented as mean +/− SEM. * p<0.0001, ** p<0.002

Entries into the center region of the open field are often interpreted to reflect risk-taking or disinhibition (Colorado et al., 2006). AIE reduced the number of entries into the center portion of the open field (Main effect of Exposure: F_(1,28)_ = 14.64, *p=0.0007*) and trend for Exposure by Sex interaction: F_(1,28)_ = 3.13, *p=0.08*) (**Fig. 3A**). Although AIE diminished the number of entries into the center portion of the arena, it did not significantly alter the amount of time spent (**Fig. 3B**), nor the total distance moved within the center portion of the field (**Fig. 3C**).

**Figure 3:**
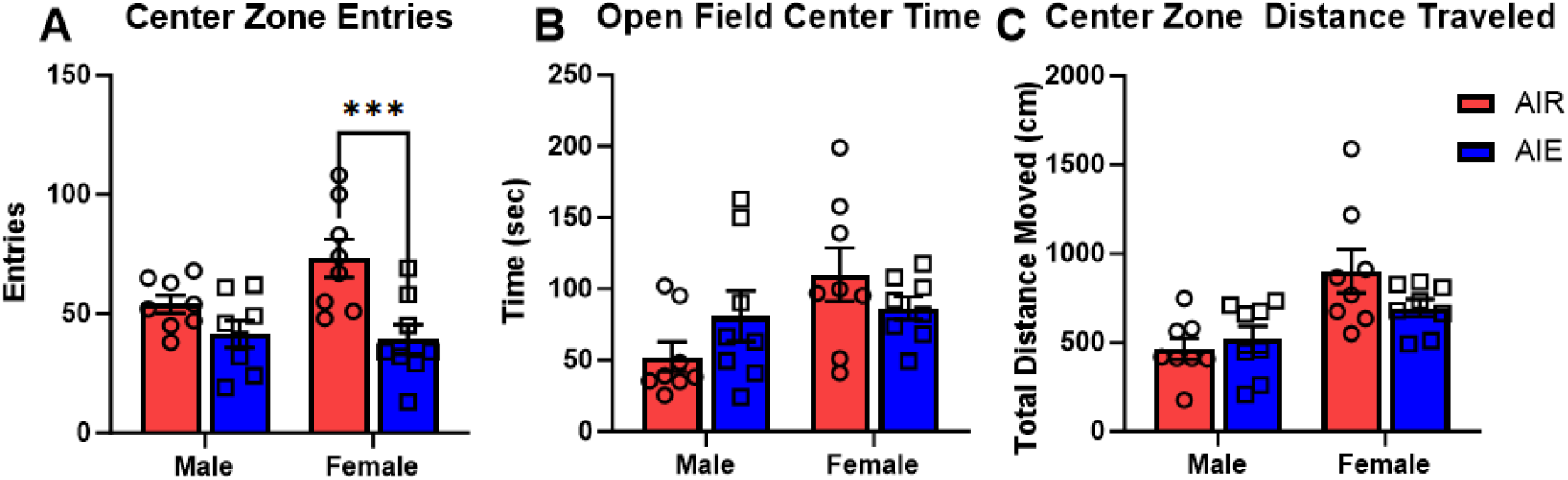
Center Zone Data. A) Entries into Center Zone, B) Time spent in Center Zone (sec), C) Distance traveled in center zone (cm). Data are presented as mean +/− SEM. * p<0.001

#### 3.2.2 Light-Dark Test

The light-dark test is used as a measure of anxiety and stress reactivity. Neither AIE nor sex altered the latency to enter the lighted area (**Fig. 4A**) [Sex (F_(1, 28)_ = 0.4649, *p=0.5010*); Exposure (F_(1, 28)_ = 0.6670, *p=0.4210*), Sex by Exposure (F(1, 28) = 0.4282, p=0.5182)], the average time spent in the lighted area (**Fig. 4B**) [Sex (F_(1, 28)_ = 0.0212, *p=0.8851*); Exposure (F_(1, 28)_ = 0.8020, *p=0.3781*); Sex by Exposure (F_(1, 28)_ = 0.0056, *p=0.9411*)], nor the average number of entries into the lighted area (**Fig. 4C**) [Sex (F_(1, 28)_ = 3.308, *p=0.0755*); Exposure (F_(1, 28)_ = 2.897, *p=0.0999*); Sex by Exposure (F_(1, 28)_ = 0.1631, *p=0.6894*)].

**Figure 4:**
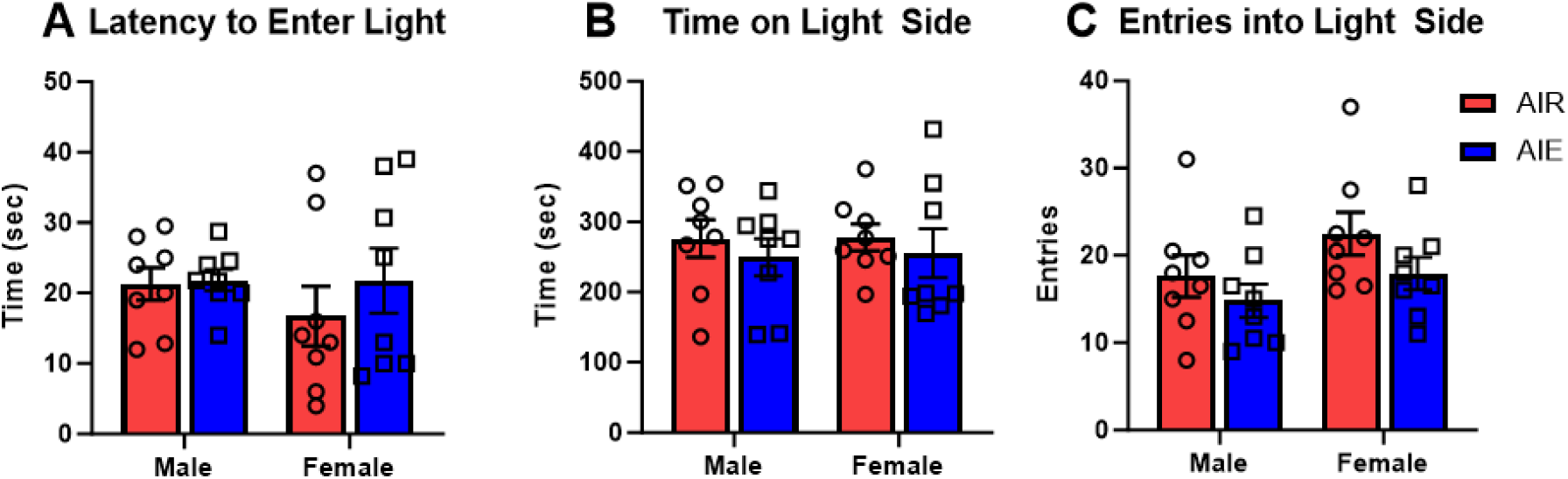
Light/Dark Test Data. A) Latency to enter light side (sec), B) Time on the light side (sec), C) Entries into the light side. Data are presented as mean +/− SEM.

#### 3.2.3 Tail Suspension Test

The TST is used as a measure of depression-like behavior and learned helplessness in mice (Cryan et al., 2005). Neither AIE (F_(1, 28)_ = 0.2250, *p=0.6390*) nor sex (F_(1, 28)_ = 2.215, *p=0.1478*) altered the latency to immobility in the test (**Fig. 5A**). However, there was a trend for immobility times to increase among females that were exposed to AIE compared to controls (*p=0.07*), but no difference in males (*p=0.25*) as supported by a significant Exposure by Sex interaction (F_(1,28)_ = 4.59, *p=0.0408*) (**Fig. 5B**). Female controls were immobile sooner than control males (p=0.0369).

**Figure 5:**
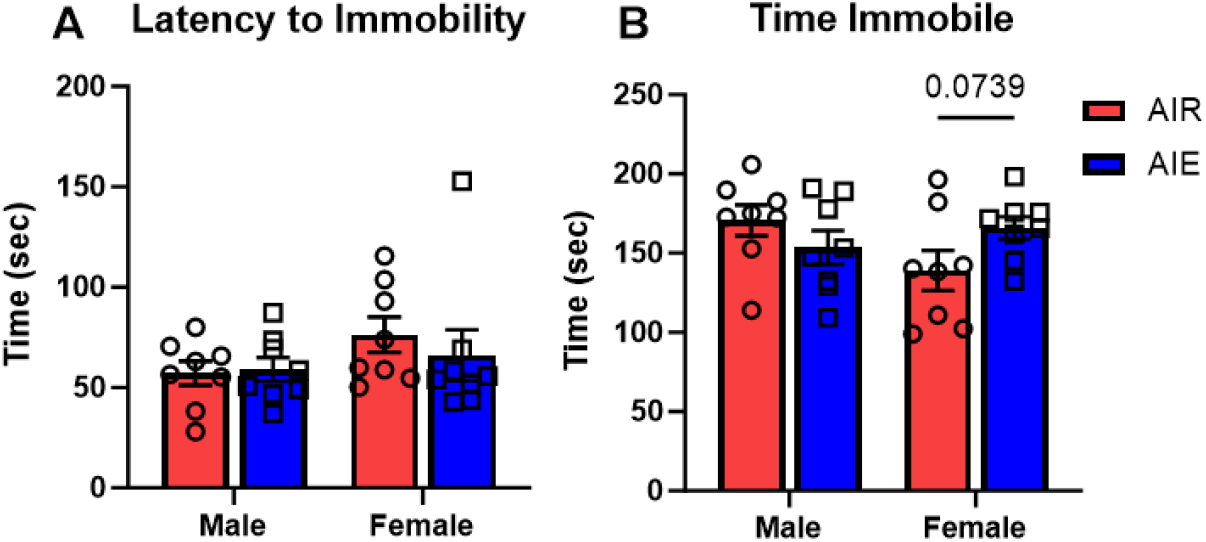
Tail Suspension Test Data. A) Latency to become immobile (sec), B) Time spent immobile (sec). Data are presented as mean +/− SEM.

#### 3.2.4 Forced Swim Stress Test

As used in this study, the FST is used as an index of stress reactivity in rodents (Commons et al., 2017; Maldonado-Devincci et al., 2014a), with time to immobility used as the dependent variable of interest. AIE exposure reduced time to immobility in males (*p=0.01*), but not females (*p=0.99*) (**Fig. 6A**) as supported by an Exposure by Sex interaction (F_(1,28)_ = 4.67, *p=0.039*) and main effect of Exposure (F_(1,28)_ = 4.52, *p=0.04*). There were no differences in time spent struggling (**Fig. 6B**) (Exposure by Sex F(_1, 28_) = 1.54, *p=0.3366*) .

**Figure 6:**
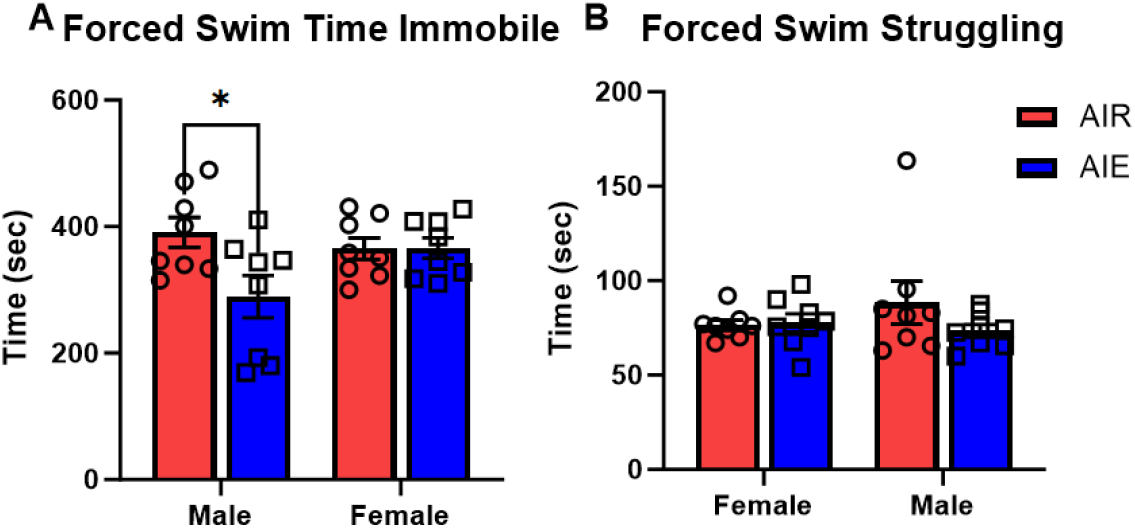
Forced Swim Stress Test Data A) Time spent immobile (sec), B) Time spent struggling (sec). Data are presented as mean +/− SEM. * p<0.05.

### 3.3 Gene Expression

We measured mRNA levels of *FMR1* and other interacting gene expressions such as *Grin2a*, *Grin2B*, *Grm5*, *PSD-95*, and *Eaat1* in the cerebellum of control and adult AIE-exposed male and female mice. We observed that mRNA levels of all these genes are not altered by AIE except *Grin2B* mRNA levels (**Fig. 7**). Interestingly, *Grin2B* mRNA expression was significantly increased by AIE in whole cerebellar tissue from both male (*p=0.033*, t-test) and female *(p=0.039*, t-test) mice. All statistical analyses for each target are shown in **Supplemental Table 2**.

**Figure 7:**
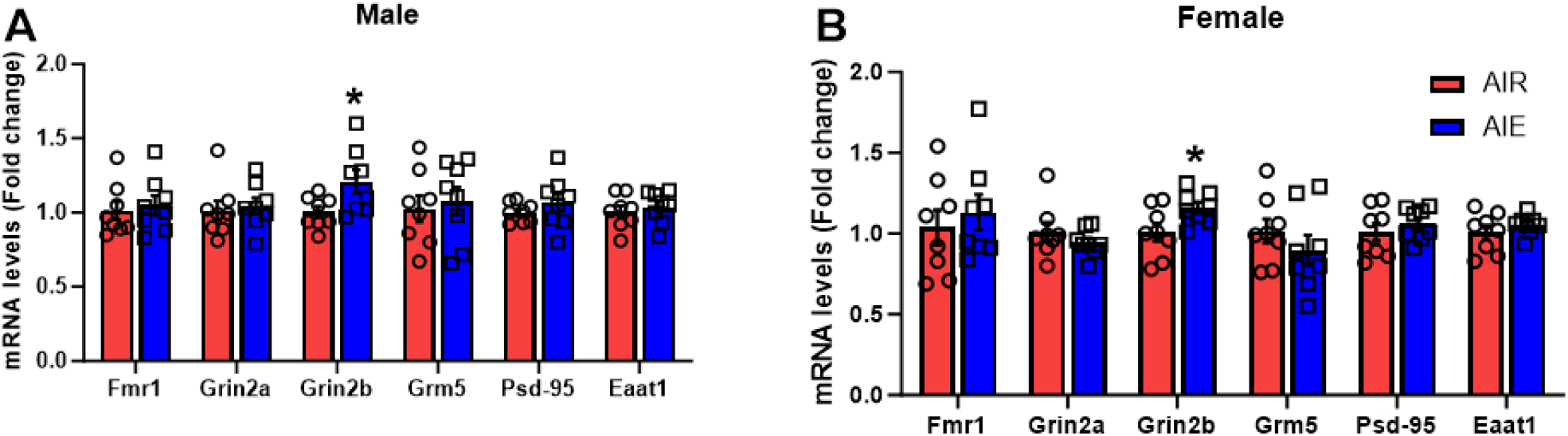
Effects of AIE on mRNA levels in the cerebellum. A) Male, B) Female cerebellar mRNA expression of FMR1 and other interacting genes. Data are presented as mean +/− SEM. * p<0.05

## 4. Discussion

The results of this study support the growing understanding that AIE alters exploratory and affective-like behaviors in adult rodents, and that some of those changes are only observed in one sex. Specifically, we found that AIE reduced open field activity in adult mice of both sexes. It also reduced exploratory rearing and entries into the center portion of the open field, though those effects were driven largely by AIE effects in females. In addition, there was a significant interaction of the effects of age and sex on behavior in the FST, indicating decreased immobility among males and suggesting that AIE exposure may lead to differential stress responsiveness between the sexes in adulthood. This interpretation is consistent with our recent finding that AIE elevated the stress responsivity of male (but not female) rats to restraint stress (Healey et al., 2023). Thus, it may be that AIE-exposed females show an altered anxiety-like/risk-taking phenotype (OFT results; center zone entries) and AIE-exposed males show an altered stress-reactivity (FST results) phenotype in adulthood. AIE had no main effect on behavior in the light-dark or tail-suspension tests.

It is notable that most of the work on AIE in rodent models has been done in rats (see (Crews et al., 2019; Robinson et al., 2021; Spear and Swartzwelder, 2014), though studies in mice have been proliferating of late (Carzoli et al., 2019; Kasten et al., 2020; Khan et al., 2023; Maldonado-Devincci et al., 2022; Piekarski et al., 2022). Studies of AIE using mice are needed because mouse models readily enable studies of genetic influence via transgenic models and inbred strains not available in rats, and because mice consume ethanol in free-choice paradigms at levels that significantly exceed free-choice self-administration in rats (Griffin, 2014; Maldonado-Devincci et al., 2022). Thus, mice provide an AIE model that has advantages for both genetic and self-administration studies. Rodent models of adolescent ethanol exposure provide translationally-relevant behavioral assays in multiple domains, a distinct peri adolescent period, and multiple brain structures that, because of their physiological consistency with parallel human brain structures, represent clinically and translationally meaningful models.

Previous literature is mixed regarding intermittent ethanol altering anxiety-like phenotypes through increases, decreases, or no change in distance traveled (Kraeuter et al., 2019; Neira et al., 2022; Quijano Cardé and De Biasi, 2022; Seemiller et al., 2022). In the present work, thigmotaxis in the open field was manifested as a decrease in movement in AIE-exposed mice in the OFT, an effect that was confined to the peripheral zone (data not shown). Future work should assess other behavioral measures including freezing episodes in both zones to determine more nuanced differences in anxiety-like behaviors in male and female mice. Similarly, we found no AIE effects on the behavior of mice in the light-dark test, while Desikan and colleagues (Desikan et al., 2014) reported that AIE-exposed male rats returned to the light portion of the testing apparatus sooner than did water-exposed male rats, and engaged in more rearing in the lighted chamber. In addition, whereas we found that AIE decreased immobility time in the FST among male mice, the previous study reported that AIE caused an increase in forced swim immobility in male rats (Desikan et al., 2014). The forced swim parameters were different between the two studies, but the observed behavioral differences suggest possible species-specific differences in stress responsivity in adulthood after AIE. Similarly, the open field and light-dark test data suggest an AIE-induced increase in risk tolerance in rats, but not in mice. These differences have methodological implications for future studies, and the species differences in enduring AIE responsivity may also lead to new mechanistic hypotheses. However, other studies have shown consistent findings of AIE effects of anxiety-like behaviors (elevated plus maze and light-dark box exploration tests) in male rats (Bohnsack et al., 2022; Kyzar et al., 2019; Pandey et al., 2015). In these studies animals were subjected to only one behavior test after AIE in adulthood.

The present study was not designed to directly assess the impact of cerebellar mechanisms on AIE-induced behavioral changes. However, the well-known relationship of cerebellar function to affective behavior (see (Schmahmann and Sherman, 1998), the effects of AIE on cerebellar structure in both humans and animal models (Coleman et al., 2014; da Silva et al., 2018; Kekkonen et al., 2021; Lisdahl et al., 2013), and the reciprocal and behaviorally relevant interactions of the cerebellum with the hippocampal formation, which is markedly modified by AIE (see (Crews et al., 2019; Robinson et al., 2021; Spear and Swartzwelder, 2014) for reviews), indicate the need to begin to address the effects of AIE on the cerebellum. Therefore, the present findings demonstrate AIE-induced increase of cerebellar *Grin2B* mRNA levels in both sexes, in the absence of changes in a number of other markers for excitatory synapse-related gene expression. This provides a specific gene expression focus for future studies of AIE on cerebellar function and a possible mechanism underlying those changes. It also suggests a possible mechanism underlying the neurotoxic effects of AIE because Grin2B codes for the GluN2B subtype of NMDA receptors, which have been associated with excitatory neurotoxicity (Zhou et al., 2013). Thus, *Grin2B* hyperexpression after AIE could relate to the previously reported cerebellar cell loss after chronic ethanol exposure (da Silva et al., 2018) through an excitotoxic mechanism because increased GluN2b synaptic drive has been shown in hippocampal circuits after AIE (Swartzwelder et al., 2017) and is associated with potentially neurotoxic extra-synaptic glutamatergic drive (Zhou et al., 2013). Thus, the present findings indicate that excitatory synaptic mechanisms in the cerebellum are altered by AIE and support the hypothesis that AIE may alter cerebellum-hippocampal interactions through those mechanisms. It is also notable that AIE did not alter the expression of *FMR1* in the cerebellum in this study, though *FMR1* expression was increased in the hippocampus after AIE (Mulholland et al., 2018), suggesting regional specificity with respect to the effects of AIE on synapse-related gene expression.

### 4.1 Limitations

Some of the limitations in this study include the age of AIE exposure was limited to only early adolescence and did not include adult comparisons. It is well known that adolescence can be divided into different developmental periods and exposure during early compared to late adolescence can induce different phenotypes including anxiety-like behavior, reward devaluation, conditioned taste aversion, peptide signaling, dopamine transmission, and ethanol sensitivity (Desikan et al., 2014; Healey et al., 2020; Saalfield and Spear, 2015; Spodnick et al., 2020; Towner and Spear, 2021; Varlinskaya et al., 2020, 2014). Multiple withdrawal time points were not used in the present study; and different behavioral and cerebellar findings may be observed after shorter or longer withdrawal time periods (Maldonado-Devincci et al., 2022). Shipping may blunt overall behavioral responding in adolescent male and female mice (Laroche et al., 2009). However, all mice were given at least a week to habituate to the animal housing facility before experimental manipulations, handled and manipulated similarly, and behavioral testing did not occur until well into adulthood. All of these limitations can be addressed in future work.

Testing order was not counterbalanced across animals and may have influenced behavioral outcomes. It is possible that previous behavioral testing could have influenced the outcomes on the latter behavioral tests, though the tests were intentionally ordered so that the most stressful tests occurred last in the behavioral testing sequence, and we did observe AIE effects on the last test given (FST). Therefore, the lack of effect we observed on the light/dark box and tail suspension tests should be interpreted cautiously. It is also possible that the behavioral testing sequence could have diminished the likelihood of observing AIE-induced changes in the expression of some of the genes assessed. Still, our goal was to assess gene expression in behaviorally tested animals, so we anticipated that limitation. Either way, it underscores the significance of the AIE effect on *Grin2B* expression in the cerebellum of both males and females.

### 4.2 Final Conclusions

These findings indicate that early AIE exposure induces long-lasting behavioral (reduced OFT distance traveled in both sexes, reduced OFT rearing in female mice, reduced OFT center zone entries in female mice, and reduced FST time immobile in male mice) and cerebellar alterations in both adult male and female mice. These findings suggest that male mice may manifest post-AIE behavioral change more in the domain of stress reactivity. In addition, the findings show, for the first time, that AIE increases the mRNA expression of *Grin2B* in the adult cerebellum. From the standpoint of translation and clinical relevance, these findings suggest important sex differences in the behavioral effects of AIE in adulthood related to anxiety and stress reactivity and point toward a specific gene expression and receptor level mechanism that could underlie those effects, thereby helping to guide the development of ameliorative treatments for the enduring effects of adolescent alcohol exposure.

## Contributors

RCW, SGK, GMW, NIH, BNJ, and MJS collected the data. AMD designed the experiments. KH, GMW, SCP, HSS, and AMD analyzed the data. KH, JGM, GMW, SCP, HSS, and AMD drafted and edited the manuscript.

## Data Availability Statement

The data that support the findings of this study are available from the corresponding author upon reasonable request.

## Acknowledgements

The authors thank Janae Baker, Kristoni Barnes, Taylor Costen, Chrystal Davis, Destinee Hamilton, Myracle Jones, Anjali Kumari, Jalen Matthews, Quamya Oglesby, Aspen Pullum, and Victoria Robinson for their expert technical assistance.

## Supplemental Tables

**Supplemental Table 1:** Behavior Testing Statistics.

|  | **F(interaction)** | **n** | **p** | **F(Sex)** | **n** | **p** | **F(AIE)** | **n** | **p** |
| --- | --- | --- | --- | --- | --- | --- | --- | --- | --- |
| *Open Field Test* |  |  |  |  |  |  |  |  |  |
| *Total Distance Moved* | F(1,28) = 1.77 | n = 32 | p = 0.194 | F(1,28) = 0.38 | n = 32 | p = 0.542 | F(1,28) = 31.49 | n = 32 | p < 0.001 |
| *Rearing* | F(1,28) = 4.72 | n = 32 | p = 0.039 | F(1,28) = 0.02 | n = 32 | p = 0.876 | F(1,28) = 9.62 | n = 32 | p = 0.004 |
| *Center Time* | F(1,28) = 3.20 | n = 32 | p = 0.0843 | F(1,28) = 4.74 | n = 32 | p = 0.038 | F(1,28) = 0.03 | n = 32 | p = 0.866 |
| *Center Entries* | F(1,28) = 3.13 | n = 32 | p = 0.088 | F(1,28) = 1.96 | n = 32 | p = 0.173 | F(1,28) = 14.64 | n = 32 | p < 0.001 |
| *Center Total Distance Moved* | F(1,28) = 2.61 | n = 32 | p = 0.117 | F(1,28) = 14.32 | n = 32 | p < 0.001 | F(1,28) = 0.888 | n = 32 | p = 0.354 |
| *Light-Dark Box* |  |  |  |  |  |  |  |  |  |
| *Latency to Enter Light Side* | F(1,28) = 0.43 | n = 32 | p = 0.518 | F(1,28) = 0.46 | n = 32 | p = 0.501 | F(1,28) = 0.667 | n = 32 | p = 0.421 |
| *Avg Time in Light Side* | F(1,28) = 0.01 | n = 32 | p = 0.941 | F(1,28) = 0.02 | n = 32 | p = 0.885 | F(1,28) = 0.80 | n = 32 | p = 0.378 |
| *Avg Entries in Light Side* | F(1,28) = 0.16 | n = 32 | p = 0.689 | F(1,28) = 3.41 | n = 32 | p = 0.076 | F(1,28) = 2.90 | n = 32 | p = 0.100 |
| *Tail Suspension Test* |  |  |  |  |  |  |  |  |  |
| *Latency to Immobility* | F(1,28) = 0.47 | n = 32 | p = 0.500 | F(1,28) = 2.22 | n = 32 | p = 0.148 | F(1,28) = 0.23 | n = 32 | p = 0.639 |
| *Time Immobile* | F(1,28) = 4.60 | n = 32 | p = 0.041 | F(1,28) = 0.91 | n = 32 | p = 0.348 | F(1,28) = 0.23 | n = 32 | p = 0.634 |
| *Forced Swim Test* |  |  |  |  |  |  |  |  |  |
| *Time Immobile* | F(1,28) = 4.67 | n = 32 | p = 0.039 | F(1,28) = 1.16 | n = 32 | p = 0.290 | F(1,28) = 4.52 | n = 32 | p = 0.042 |
| *Struggle Behavior* | F(1,28) = 1.54 | n = 32 | p = 0.226 | F(1,28) = 0.347 | n = 32 | p = 0.561 | F(1,28) = 1.10 | n = 32 | p = 0.304 |

**Supplemental Table 2:** mRNA Statistics.

| **Target** | **Male** |  |  | **Female** |  |  |
| --- | --- | --- | --- | --- | --- | --- |
|  | (AIE) | n | p | (AIE) | n | p |
| ***Fmr1*** | t(14) = 0.40 | 16 | p = 0.694 | t(14) = 0.62 | 16 | p = 0.545 |
| ***Grin2a*** | t(14) = 0.38 | 16 | p = 0.707 | U(n1, n2 = 8) = 22 | 16 | p = 0.314 |
| ***Grin2b*** | t(14) = 2.39 | 16 | p = 0.031 | t(14) = 2.27 | 16 | p = 0.039 |
| ***Grm5*** | t(14) = 0.40 | 16 | p = 0.696 | t(14) = 1.01 | 16 | p = 0.330 |
| ***pSD-95*** | t(14) = 1.03 | 16 | p = 0.322 | t(14) = 0.79 | 16 | p = 0.444 |
| ***Eaat1*** | t(14) = 0.54 | 16 | p = 0.600 | t(14) = 0.95 | 16 | p = 0.360 |

